# Opportunities and pitfalls in preclinical cerebral blood flow mapping using arterial spin labelling MRI: insights from multicentre data

**DOI:** 10.64898/2026.06.22.733736

**Authors:** Sara P. Monteiro, Dylan Dunkwu, Steven Reynolds, Patrícia Figueiredo, Noam Shemesh, Yolanda Ohene, Isabel N. Christie

## Abstract

Cerebral blood flow (CBF) is a quantitative metric for mapping perfusion. While the prototypical MRI approach arterial spin labelling (ASL) is well-validated in humans, the reproducibility of rodent ASL mapping remains poor, limiting translational impact. To address this gap, we used both newly acquired and analysis of previously published data to illustrate biological and physical sources of variation in CBF measured with ASL. Via a meta-analysis, we quantified the variation in CBF reported from the cortex of healthy rodents. A total of 23 mouse studies (343 data points) and 5 rat studies (41 data points) met the inclusion criteria. We demonstrate that reported CBF values exhibit a broad variability (50-400 ml/100g/min) driven primarily by experimental confounds rather than physiological differences. Our meta-analysis explores which factors cause variance in perfusion rates measured. Our experimental data highlight biological factors, particularly the choice of anaesthesia (e.g., isoflurane vs. medetomidine) and strain variations, that alter baseline CBF. Our work, reflecting both state-of-the-art and conventional practice in preclinical imaging, highlights the need to account for multiple sources of variability. Establishing community guidelines for rigorous ASL calibration and physiological monitoring will support improved study design and accelerate translational alignment between rodent and human perfusion measurements.

**Graphical abstract:** 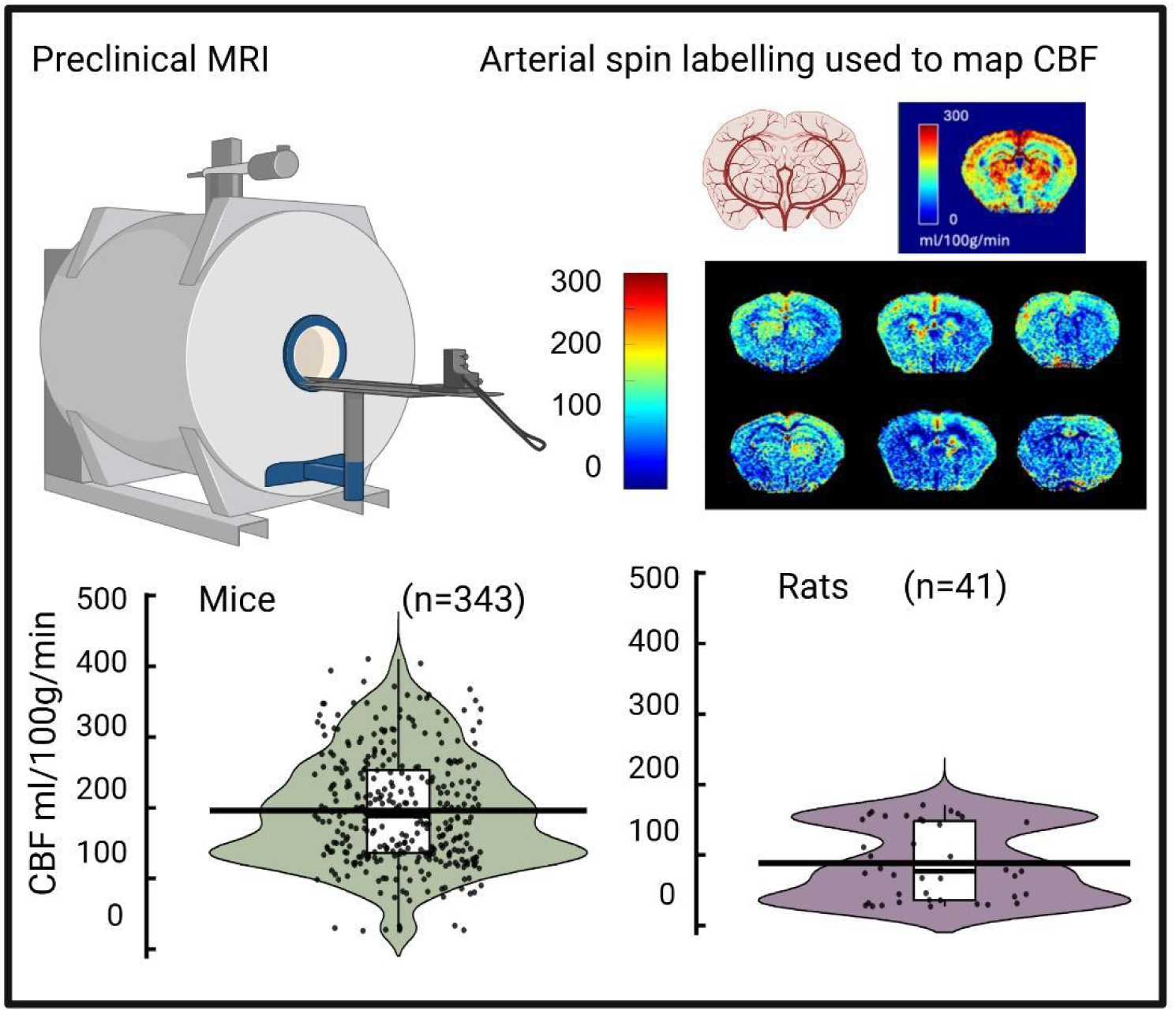

## Introduction

Cerebral blood flow (CBF) is a fundamental physiological parameter that ensures the continuous exchange of metabolic gases (oxygen and carbon dioxide) and the supply of glucose to the brain (1, 2). Since cerebral perfusion rate reflects dynamic homeostatic mechanisms, CBF has emerged as a critical and sensitive indicator of brain health and disease. Arterial spin labelling (ASL) MRI is a powerful tool for quantifying CBF (3–5). ASL uses radiofrequency pulses to magnetically label endogenous blood water in the arteries, and a series of images are acquired to capture the passage of the labelled blood through the circulation and its exchange into the brain tissue (6). Since the technique does not require an exogenous contrast, it is accessible to many populations, including children and people with poor kidney function, and has been applied widely in human research.

The successful translation of ASL into clinical practice has been supported by a concerted effort to harmonise acquisition and analysis protocols (7–16). Comprehensive community guidelines and large-scale multi-centre studies have been conducted to demonstrate reproducibility of human CBF measurements (13–22), ensuring they remain robust against variations in hardware vendor or field strength. By contrast with human ASL MRI, such a high level of standardisation has yet to be achieved in the preclinical domain. Rodent models, particularly transgenic mice, engineered to develop specific pathophysiology, and in which invasive methods can be used for validation, are indispensable for understanding mechanisms of disease in general and vascular dysfunction in particular (17, 18). Yet, the preclinical literature is characterised by a high degree of variability in reported CBF values, which are influenced by a disparate range of experimental configurations rather than intrinsic biological differences.

The specific pulse sequences used to acquire data and label endogenous blood water have been a source of methodological variability in preclinical CBF estimates. Several ASL techniques are typically implemented in rodent studies. Figure 1 presents the general principles of ASL, highlighting the two most common labelling strategies. Pulsed ASL (PASL) uses a single short RF pulse to label a slab of blood symmetrically about the imaging slice (19). The most common PASL sequence is flow-sensitive alternating inversion recovery (FAIR), which requires the collection of two inversion recovery images: one slice-selective and one non-slice-selective (Figure 1Ai). FAIR has been widely used due to its relative technical simplicity and robustness to B0 inhomogeneities. However, the field is increasingly transitioning toward pseudo-continuous ASL (pCASL) to improve labelling efficiency and better model the labelled blood as a bolus (20–23). Briefly, pCASL uses a long train of short RF pulses to mimic continuous ASL. The time between labelling and acquisition is referred to as the post-labelling delay (PLD) (Figure 1Aii).

**Figure 1.**
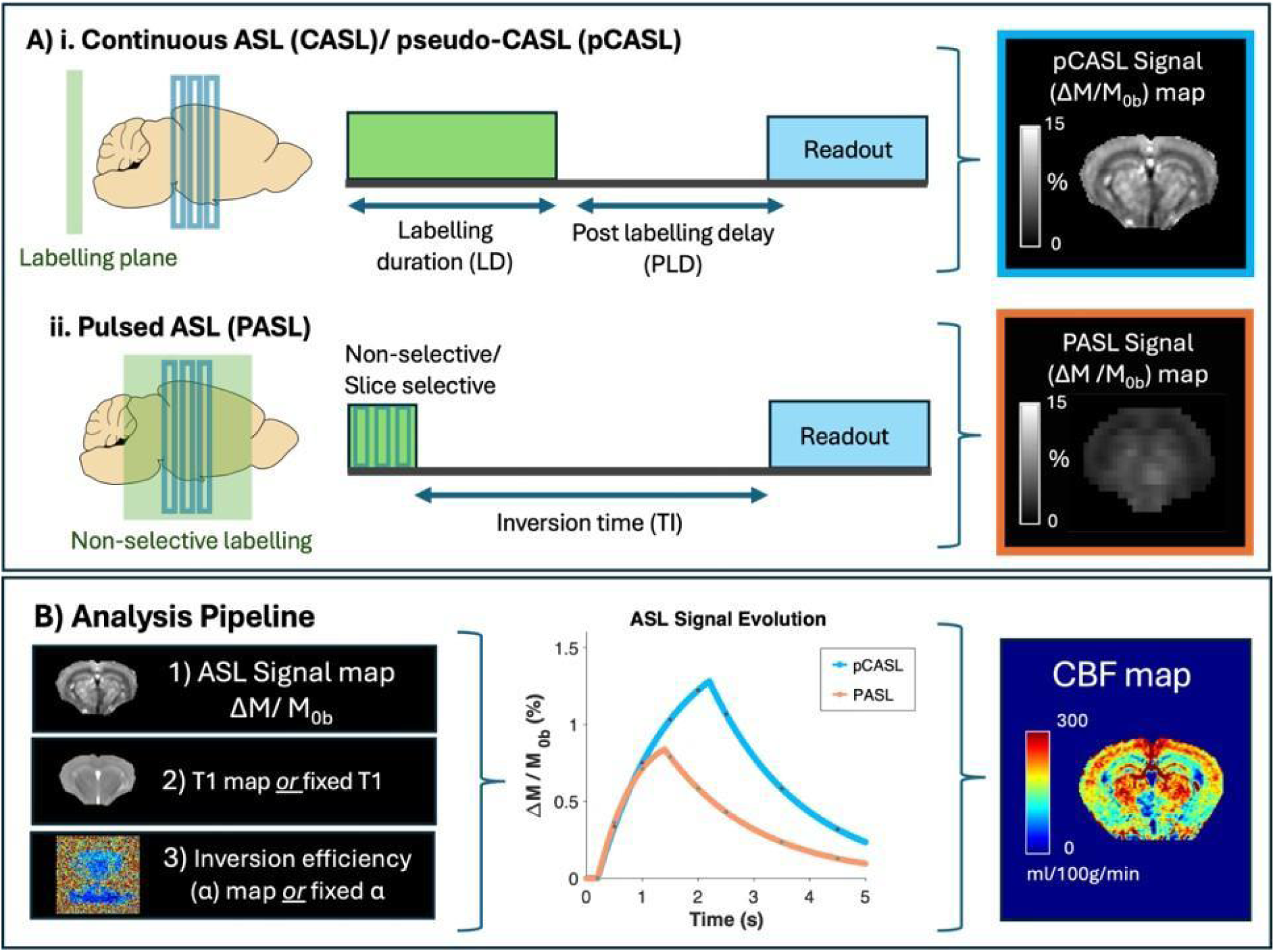
**General overview of arterial spin labelling (ASL) in rodents, including the main acquisition and analysis methods**. ***A*** Pulse sequence diagram with example ASL Signal (ΔM) maps for i. Pulsed ASL (PASL) ii. Pseudo-continuous ASL (pCASL). ***B*** Analysis pipeline using the Buxton model to fit the ASL control-label ΔM signal map, T1 map or fixed T1, and inversion efficiency map or fixed inversion efficiency, to derive a quantitative CBF map (ml/100g/min).

For CBF quantification in both PASL and pCASL approaches, a general kinetic model (e.g., the Buxton model (6) is typically used to fit the ASL signal from the control-label images. CBF (and arterial transit time) can be estimated by acquiring multiple ASL images at increasing delays, inversion time (TI) or post labelling delay (PLD), between adiabatic inversion and image acquisition. In the PASL FAIR method two images are acquired with either selective or non-selective adiabatic inversion. Whereas in pCASL labelling is performed in the carotid arteries. As outlined in Figure 1B, the CBF calculation requires incorporating T1 values (either from a measured map or an assumed fixed T1 and inversion efficiency values (measured or assumed). Despite extensive use of ASL in rodents, quantification challenges remain, such as the vulnerability of labelling efficiency to complex vascular geometry and the necessity of mapping precise arterial transit times (21).

Accurately capturing these fine cerebrovascular dynamics in the small rodent brain requires maximising the signal-to-noise ratio (SNR) to achieve high-spatial-resolution CBF maps. This challenge is typically met by employing ultra-high field (UHF) magnetic field strengths (3, 23, 24) and signal boosting hardware, such as cryogenic coils (25). However, these hardware configurations vary more widely between preclinical sites than in clinical environments. Furthermore, higher field strengths can exacerbate B0 and B1 inhomogeneities at the labelling plane, compounding the sequence-related quantification issues.

Finally, there are the additional physiological challenges of rodent imaging. While some recent studies perform imaging in awake animals to capture true baseline physiology, the vast majority of preclinical ASL requires anaesthesia or sedation to prevent motion (26). Although breathing rate and body temperature are closely monitored, the choice and depth of sedatives perturb respiratory rates, blood gases, and mean arterial blood pressure, introducing an expansive range of biological variability into preclinical studies (27, 28).

Altogether, there is a growing need to characterise the variability and identify its sources, in this way gaining opportunities for more reliable and reproducible quantification of preclinical CBF. This study set out to define a normative physiological range for healthy adult rodent CBF measured via ASL under anaesthesia. First, using *de novo* multi-site experiments, we illustrate how biological factors drive perfusion differences both within and across laboratories. Second, we perform a first of its kind meta-analysis of the published literature to quantify the impact of methodological choices on reported CBF values.

## Methods

Our work includes two main directions (1) a study showcasing CBF variability within a single site (shown in two separate sites with two distinct protocols), and (2) a meta-analysis to quantify and dissect various factors leading to variability across different sites.

### New data acquired in mice at two sites

MRI experiments were performed in two sites using two different systems located in two separate countries (UK and Portugal).

### Site 1 (UK)

Experiments were performed on male mice in accordance with the European Commission Directive 86/609/EEC (European Convention for the Protection of Vertebrate Animals used for Experimental and Other Scientific Purposes) and the United Kingdom Home Office (Scientific Procedures) Act (1986) with project approval from the Institutional Animal Care and Use Committee. The results of the animal experimentations are reported in accordance with ARRIVE guidelines.

**Animal Preparation:** Adult male C57BL/g mice were used (weights 20-25 g, *n* = 7). All animals were sedated with isoflurane (4%), followed by subcutaneous medetomidine bolus (0.4 mg/kg, and infusion 0.8 mg/kg/hr). Isoflurane was discontinued during cerebrovascular imaging. The free breathing mice were supplied with supplemental oxygen via a nosecone and gas challenges were delivered via the nosecone. Body temperature was maintained at 37°C and respiratory rate was maintained between 60-90 breaths per minute.

**MRI experiments:** All MRI experiments were performed using a 9.4T MRI scanner (Agilent Inc.), a 72 mm inner diameter volume coil for radio frequency transmission (Rapid Biomedical) and a 2-channel array surface coil (Rapid Biomedical) for signal reception. A PASL sequence, flow sensitive alternating inversion recovery with echo planar imaging readout (FAIR-EPI) was used to acquire the data (EPI readout: TE = 5.8 ms, TR = 5 s, TI = 2 s, 3 contiguous slices, 1 mm slice thickness, with matrix size = 192 x 192). To achieve higher temporal resolution (30 s per scan), the perfusion data reported were acquired with a single TI (2s). Repeated images were acquired for 10 mins at normoxia, 5 mins at hypercapnia and 3.5 mins at normoxia.

### Data Analysis Site 1 (UK)

To quantify CBF, the preliminary data were fitted to the established model (6). FAIR-EPI images were analysed using a custom Matlab script (R2025a, Mathworks, Natick, MA). The non-selective image vs TI (single TI described above) were fitted to extract a T1 value from within an ROI containing animal cerebral cortex. Selective and non-selective inversion images were then subtracted and the mean intensity with the same ROI was fitted to a general kinetic pulsed ASL perfusion model as described in (6). CBF (ml/min/100g), bolus and transit times were estimated, along with R-squared from the least-squares algorithm.

### Site 2 (Portugal)

Animal experiments were conducted according to the European Directive 2010/63 and preapproved by competent authorities.

**Animal Preparation:** Adult female C57BL/g mice (∼12 weeks old, weights 20–25g, total *n* = 9) were used for anaesthesia comparison experiment (*n* = 6) and strain comparison experiment (*n =* 3). Three adult female wildtypes of the transgenic mouse model (C57BL/6-DBA/2) were used for the strain comparison experiment (*n* = 3). For both experiments, all animals were sedated using isoflurane concentrations ranging from 1.5% to 2.5%, with the exception of three animals used for anaesthesia comparison. These mice were sedated with medetomidine: following isoflurane induction, a bolus of medetomidine solution (1:10 dilution in saline of 1 mg/ml medetomidine solution (Vetpharma Animal Health S.L., Barcelona, Spain) was administered via subcutaneous injection (0.05 mg/kg) using a syringe pump (GenieTouch, Kent Scientific, Torrington, Connecticut, USA). Ten minutes after the initial bolus, a subcutaneous constant infusion of 0.1 mg/kg/h medetomidine was initiated. Respiratory rates maintained between 60 and 90 breaths per minute.

**MRI experiments:** Experiments were conducted on a 9.4 T Bruker Biospec Scanner with an

86 mm volume coil for transmittance and a 4-element array cryogenic coil for signal reception. An unbalanced pCASL sequence was used as described in (20). The labelling plane was positioned at the mouse neck (∼8 mm below the isocenter), the labelling duration (LD) was set to 3 s followed by a 300 ms post-labelling delay (PLD). Inversion was achieved through a train of Hanning window-shaped pulses: 400 μs duration, 800 μs pulse rate, B_1_ of 5 μT, G_max_/G_ave_ of 45/5mT/m, where G_max_ is the gradient applied during the RF pulse.

For high-resolution pCASL, a single-shot EPI was implemented: FOV = 12 x 12 mm^2^, slice thickness = 0.5 mm, slice gap = 0.35 mm, matrix = 120 x 120 resulting in a spatial resolution of 100 x 100 m^2^, TR/TE = 4000/25 ms, 30 repetitions, T_acq_= 4 min. For CBF quantification, the T1 map was obtained from an inversion recovery sequence. A pCASL encoded FLASH was employed to estimate the inversion efficiency (IE) 3 mm above the labelling plane (PLD of 0 ms, LD of 200 ms) (20).

Tissue T1 mapping was performed using an inversion recovery spin-echo EPI sequence (TR/TE = 10,000/19 ms; 18 inversion times (TI) ranging from 30 to 10,000 ms; T_acq_ = 4 min) and an adiabatic full-passage shaped pulse designed via the Shinar-Le Roux algorithm (bandwidth = 5000 Hz, length = 13.56 ms).

In addition, a pCASL flow-compensated encoded fast low angle shot (fc-FLASH) sequence was used to estimate the inversion efficiency (α) by measuring the signal 3 mm above the labeling plane (1 repetition, TR/TE = 225/5.6 ms; 1 mm slice thickness, 172 X 172 µm² in-plane resolution, 2 averages, PLD = 0 ms, LD = 200 ms, T_acq_ = 3 min 30 s)

### Data Analysis Site 2 (Portugal)

#### Inversion Efficiency

For each animal and experimental configuration, the inversion efficiency (α) was derived from complex-reconstructed pCASL-encoded FLASH images. For both the left and right carotids, α was calculated using the following expression:

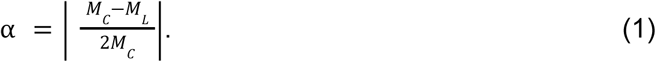

Here, *M_C_* and *M_L_* denote the complex signals from the control and label scans, respectively, averaged within manually delineated carotid regions of interest (ROIs). The final representative value of α for each subject was determined by averaging the results from both carotids.

### T1 maps

T1 maps were generated for each subject and spatial resolution by fitting the longitudinal relaxation model to voxelwise data acquired via an inversion recovery spin-echo EPI sequence. Parameter estimation was performed using the Levenberg-Marquardt algorithm based on the following equation:

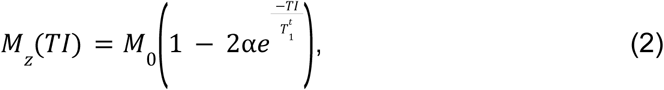

where *M_z_*(*TI*) represents the longitudinal magnetization at a specific inversion time (*TI*), *M*_0_ denotes the equilibrium magnetization, and T*^t^*_1_is the longitudinal relaxation time of the tissue.

### CBF maps

CBF maps (ml/100g/min) were calculated pixel-by-pixel (10) according to:

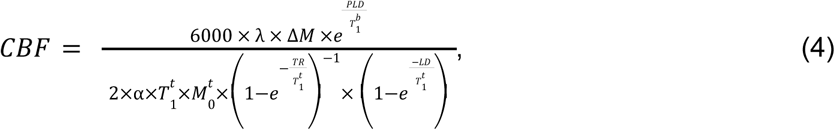

where λ is the blood-tissue partition coefficient of water (considered to be 0.9mL/g (29), and *T^b^*_1_ *T^t^*_1_ and are the longitudinal relaxation times of blood (considered to be 2430 ms at 9.4 T (30) and tissue (obtained from the voxelwise T1 map) respectively, and α is the value obtained from the inversion efficiency acquisition (Equation (1)), 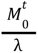 and is the equilibrium magnetisation of blood, where *M^t^*_0_is the tissue magnetisation in the average control image.

### Multi-site metadata

The meta-analysis was performed in accordance with a published protocol (31). The search terms included: perfusion, MRI, arterial spin labelling, mouse, rat, brain, ASL. The inclusion criteria were healthy wild type animals without any known health problems. The exclusion criteria were an upper age limit of 17 months, papers published before 2010, field strength below 3T, model of disease or injury, any surgical intervention such as sham surgeries. The total number of published articles which met the search criteria was 28, 23 for mice and 5 for rats. The included data are visualised in the PRISMA flow chart in Figure 2.

**Figure 2.**
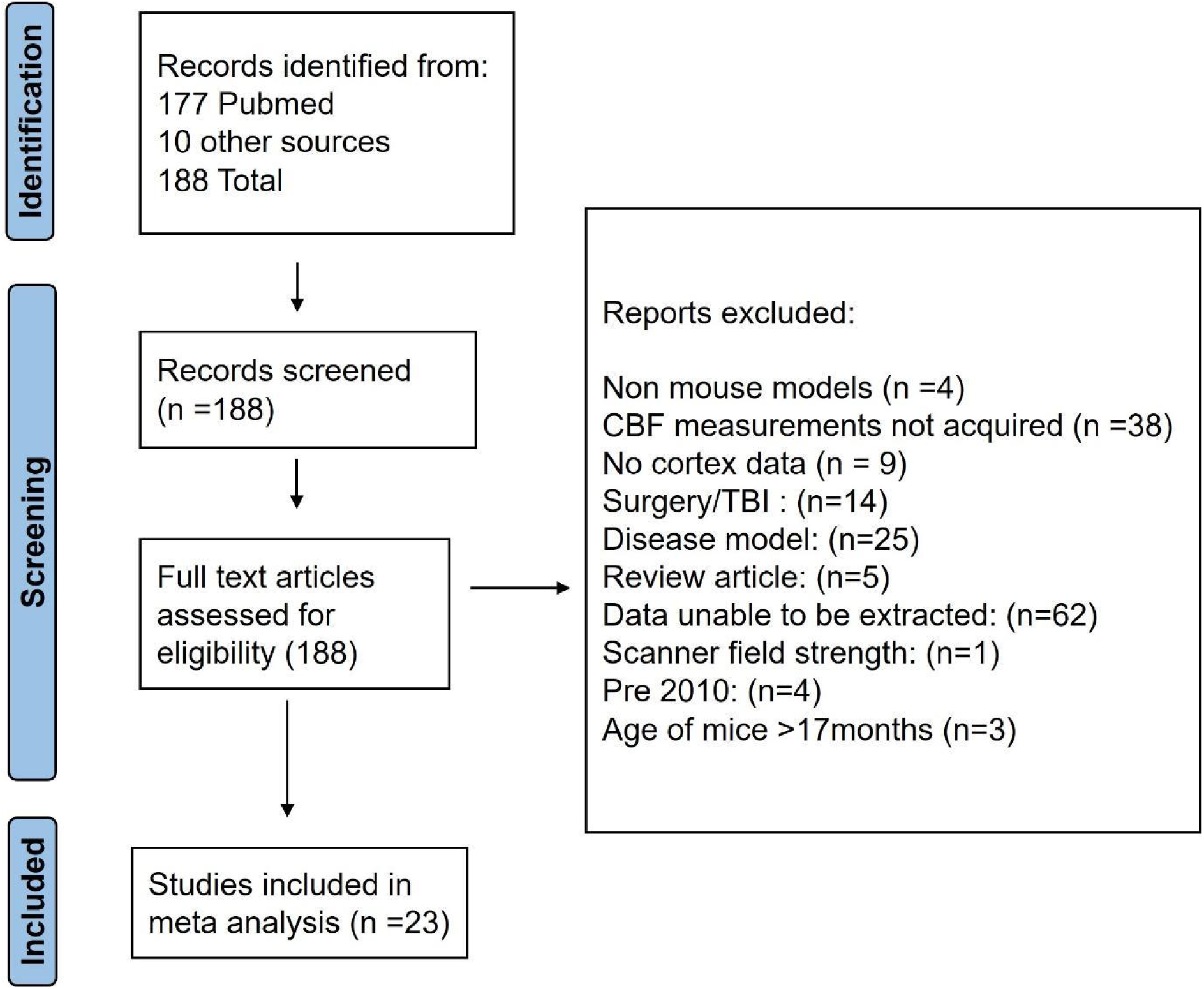
PRISMA flow diagram outlining the study selection criteria.

### Data extraction

Cortical CBF values were extracted and tabulated. If the information was not available from the full text, authors were contacted to obtain the relevant data or data were extracted using WebPlotDigitizer (https://automeris.io/).

### Metadata analysis

The data points were collated to find the mean cortical CBF values for the mouse brain and the rat brain. A Shapiro-Wilk test showed the data for mice and rats was not normally distributed. Therefore, to compare the grouped averages we conducted non-parametric Mann Whitney tests.

We sought to test whether field strength – assuming it is an independent variable – affected CBF measured, accounting for the effect of sequence, strain, anaesthesia and the covariate of age. We assumed homogeneity of the regression slope.

We then sought to test whether pulse sequence – considered an independent variable – affected CBF, accounting for the effects of field strength, strain, anaesthesia and the covariate age. We assumed homogeneity of the regression slope.

A post hoc analysis using Tukey’s HSD to complete a pairwise comparison was performed (p <0.001). To avoid the risk of interaction between field strength and sequence, PASL and pCASL data were tabulated, plotted separately and assessed with a t test.

## Results

### Changes in CBF due to varying physiology (*De Novo* Experiments)

To showcase the large spread of CBF values when physiological factors are varied, we first evaluated the impact of distinct physiological states and genetic backgrounds, in two different sites. Figure 3 summarises these comparisons.

**Figure 3.**
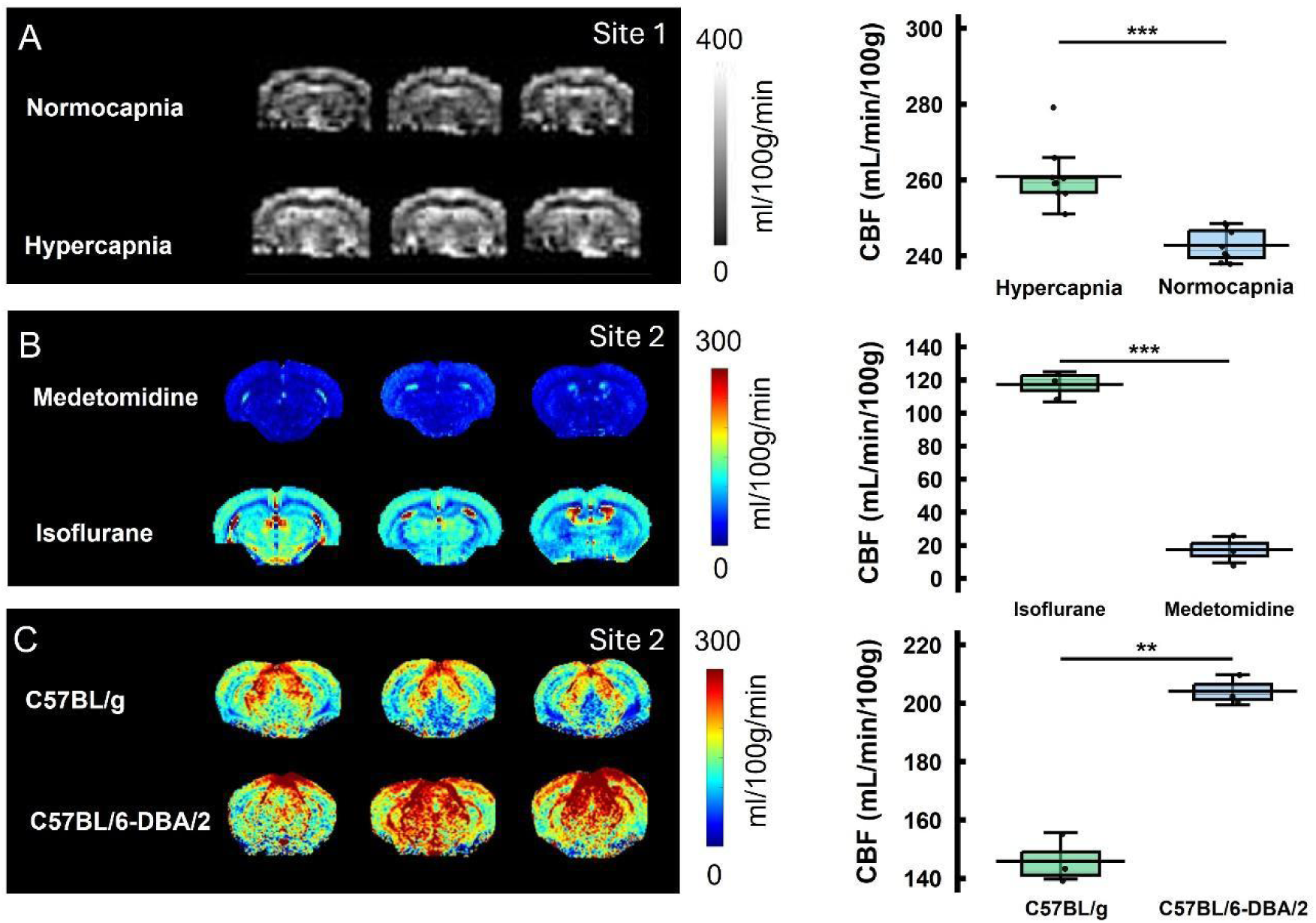
**Key biological factors that contribute to variability of ASL measurements of cerebral blood flow**; ***A*** CBF maps (left) from a representative mouse showing normocapnia and hypercapnia ROI analysis (right, *p* <0.001, *n = 7)*; ***B*** CBF maps (left) from the same animal showing the effect of anaesthetic on the rate of cerebral perfusion, ROI cortex analysis (right, strains *p* <0.001, *n = 3*) ***C*** CBF maps (left) from two different mouse strains (C57BL/g and C57BL/6-DBA/2) and ROI analysis (right, *p* <0.001, *n = 3*).

Changes in inspired CO_2_ concentrations significantly alter perfusion. Figure 3A shows a representative CBF map from a single animal in conditions of normocapnia and hypercapnia. Figure 3A (right) illustrates cortical increases in the rate of perfusion from 240 to 260 ml/100g/min when blood CO_2_ is elevated above 45 mmHg (t test p <0.001).

Figure 3B shows how anaesthetic regime can produce distinct physiological conditions that alter CBF values. Mice scanned under isoflurane anaesthesia exhibited higher brain perfusion, with mean values in the cortex in the order of 100 mL/100g/min. In contrast, mice scanned under medetomidine showed a comparatively reduced CBF, with values all remaining below 50 mL/100g/min, (t test p <0.001).

Interestingly, under identical anaesthetic and physiological conditions, we find that mouse strain is a significant biological source for CBF values (Figures 3C). Baseline perfusion differences are observed between the common wild-type strain C57BL/6 and C57BL/6-DBA/2 (right), and these differences reach statistical significance (t test p <0.001).

### Literature CBF Values (Meta-Analysis)

We next assessed the range of CBF values reported across the preclinical MRI literature in healthy adult animals. We analysed and reported baseline CBF values from 28 peer-reviewed studies - comprising 23 mouse studies (343 data points) and 5 rat studies (41 data points) - published between 2010 and 2025.

Prior to any stratification by experimental condition, we note that the reported perfusion values exhibited a broad distribution. Figure 4A shows the distribution of the 343 cortical CBF values obtained from mice, and Figure 4B displays the distribution for rats. Estimates in both species spanned a wide range from roughly 50 to 400 ml/100g/min prior to any stratification. On average, the rate of perfusion reported for rats was significantly lower than of mice (Mann Whitney p <0.001), although data are only segregated by species and not by any other factors.

**Figure 4:**
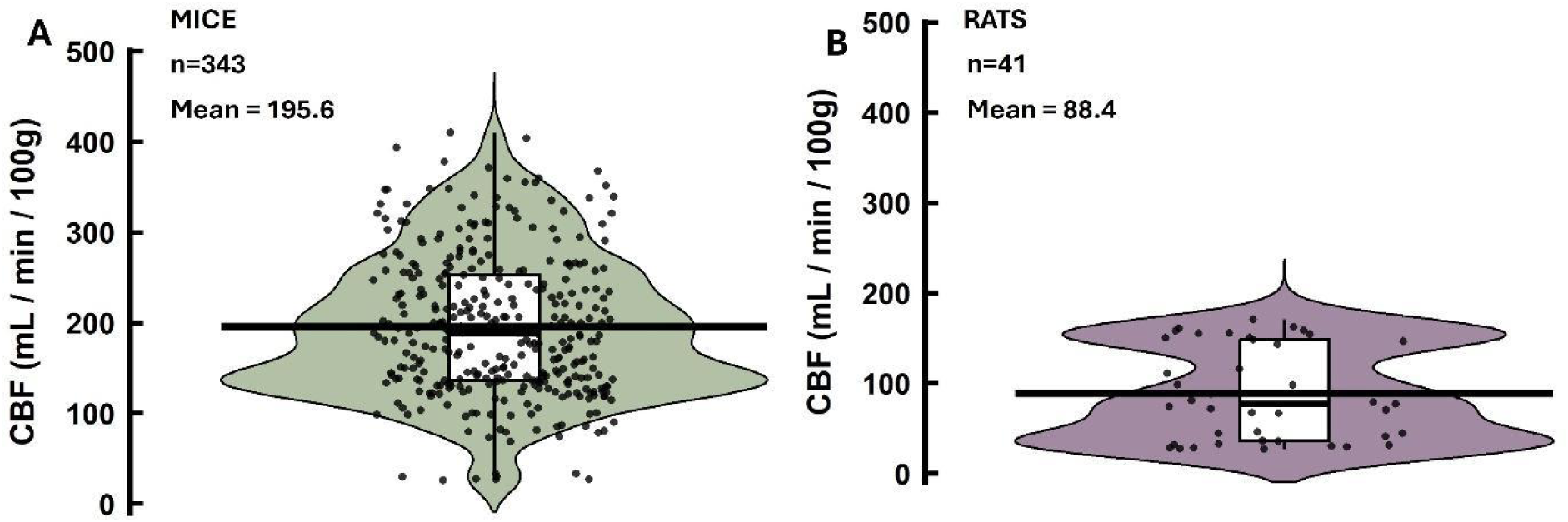
Global metadata describing the average rate of cerebral perfusion in the cortex of healthy mice and rats under anaesthesia. Mean value of 196 ml/min/100g calculated in mice and mean value of 88 ml/min/100g calculated in rats (Mann Whitney *p* <0.001).

### Methodological Drivers of CBF Variability

Given the limited number of rat studies identified (*n* = 5 studies, *n* = 41 data points), we restricted the subsequent stratification analysis exclusively to the richer mice dataset (*n* = 23 studies, *n* = 343 data points). The impact of experimental variables on reported mouse CBF is summarised in Figure 5A. Figure 5A shows perfusion values measured in mice, colour coded by site. In Figure 5B the data points are grouped by the experimental conditions field strength and sequence. The choice of labelling strategy (PASL or pCASL) appears to impact the reported physiology, as depicted in Figure 5B/C. Isolating the sequence type, and controlling for covariates age, field strength, anaesthetic, etc revealed that data acquired with a PASL strategy yielded a higher average CBF than data acquired with PASL (adjusted means are respectively: 208 ml/100g/min and pCASL = 152 ml/100g/min, p <0.01 F value 7.54). Notably, the inflationary effect of the PASL labelling strategy appeared to act independently of the field strength (Figure 5B; PASL p <0.001, pCASL p = 0.02, t-test). When plotting reported CBF as a function of magnetic field strength (Figures 5D) and controlling for the covariates of age, strain, and anaesthetic type, a methodological bias emerged. We observed that the mean CBF values measured at 9.4 T were consistently higher than those reported at 7T (p <0.001, F value 116.44, multiple linear regression followed by pairwise comparison, Figure 5D). The adjusted means for 7T = 126 ml/100g/min and 9.4T = 219 ml/100g/min.

**Figure 5:**
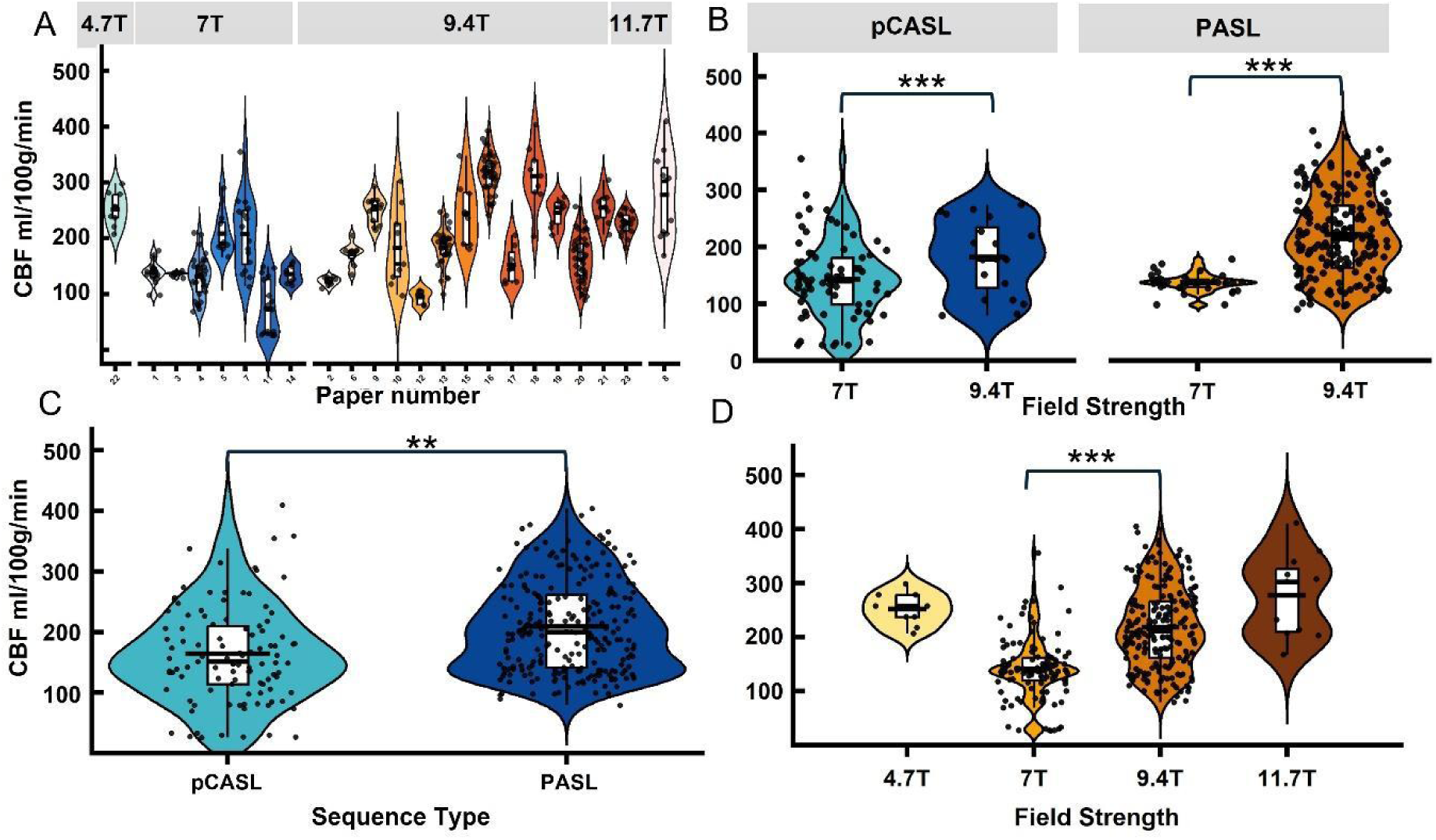
Field strength and sequence type influence average values recorded *A* Mouse cortex perfusion values separated by field strength and colour coded by site (23 in total); *B* Mouse cortex data separated by field strength and sequence choice (pulsed ASL (PASL) *p* = <0.001, pseudo-continuous ASL (pCASL) *p* = 0.024, t-test followed by pairwise comparison); *C* Mouse cortex perfusion data separated by sequence type (PASL or pCASL); *D* Mouse cortex perfusion values segregated by field strength (*p* = <0.001). Longer black line = mean average).

### Secondary Metadata Variables

Finally, we evaluated secondary covariates extracted from the metadata. Age, evaluated as a continuous variable in the multiple linear regressions was binned into three groups (2-6, 7-11, and 12-17 months) and graphed in Figure 6A. The age of the mice emergesd as a significant driver of CBF variability in the metadata when other factors were controlled, at 7-11 and 12-17 months CBF significantly increased compared to 2-6 months, although we purposefully excluded data acquired beyond 17 months. Regarding strain, we binned the data into four groups which were unbalanced by the predominance of C57 lineage, no statistical analysis was made because the groups are unbalanced. Lastly, anaesthetic regimes across studies were grouped into six categories (isoflurane <1.5%, isoflurane 1.5%, isoflurane >1.5%, MED medetomidine, isoflurane and medetomidine, urethane and α -chloralose). Anaesthesia emerged as a significant factor in the multiple linear regression (p <0.001) and post-hoc testing with Tukey’s HSD as a pairwise comparison revealed significant differences between many of the anaesthetic regimens used (Figure 6) (32, 33).

**Figure 6:**
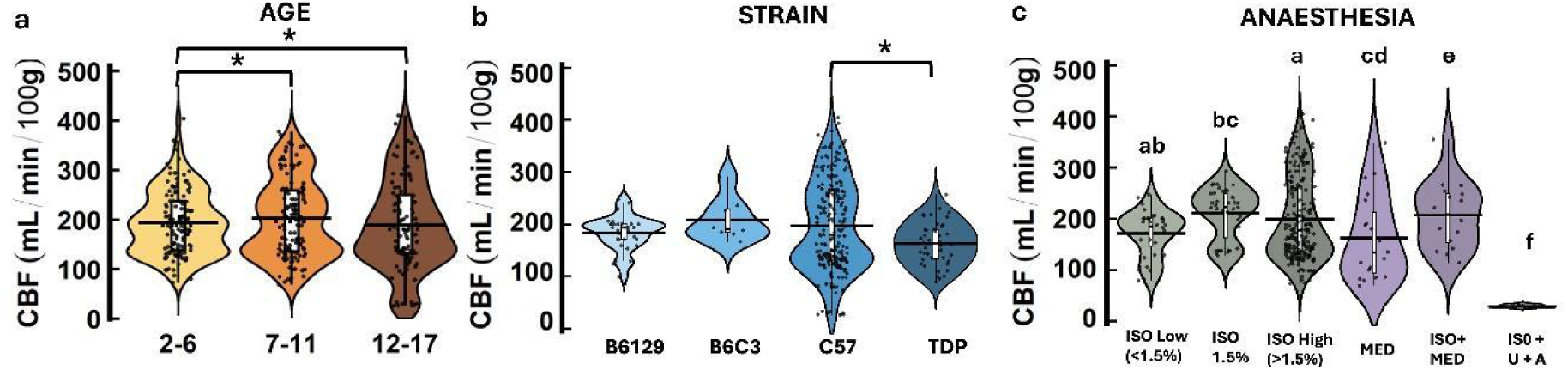
M**o**use **metadata graphed by age, strain and anaesthetic without statistical analysis *A*** A significant effect of age on mouse cortex perfusion values at 2-6 months compared to 7-11 (p =0.004) and 12-17 (p =0.004) months ; ***B*** C57 is most common strain used for CBF studies; ***C*** Cortical mouse metadata stratified by anaesthetic reported in research articles. Groups labelled with different lowercase letters are significantly different(p<0.05).

## Discussion

Our work provides the first systematic examination of the physical, biological and physiological sources of variability in preclinical CBF measurements obtained with ASL. Across the meta-analysis, the reported cortical CBF values in healthy rodents span a broad range. In mice, values from approximately 50 to 400 ml/100g/min are reported under anaesthesia, while in rats, values range from approximately 50 to 150 ml/100g/min. Such a large dispersion is difficult to reconcile with the known constraints imposed by cerebral metabolism and autoregulatory physiology (26, 34). The results presented here demonstrate that the range of reported CBF values likely does not reflect intrinsic physiological variability. Instead, this spread emerges from complex interactions between labelling strategies, magnetic field strength, anaesthetic regime, and physiological state (depressed respiratory rate, respiratory acidosis, depressed blood pressure), rather than from genuine inter-animal differences in basal perfusion.

### Sources of variability

Our meta-analysis suggests that a higher magnetic field strength (9.4T vs. 7T) systematically inflates CBF values. It is possible other unknown factors may cause this bias, since the data are pooled from many sources. A more sophisticated analysis of the data, interrogating acquisition methods and analysis protocol of all sites would be needed to verify this finding. There is no established physiological or physics mechanism by which baseline cerebral perfusion should systematically increase with magnetic field strength. Therefore, this relationship may be best understood as a methodological artifact. One possible explanation may arise from the fact that at ultra-high fields, the longitudinal relaxation time of blood markedly increases (30). Kinetic models used for ASL quantification rely on accurate values of T1b to correct for label decay (10). If field-strength-specific T1b values are not used, decay correction becomes inaccurate and leads to inflated CBF estimates. Furthermore, higher field strengths also exacerbate B0 and B1 inhomogeneities at the labelling plane (20, 21). While increased SNR at ultra-high field improves apparent image quality, it does not compensate for systematic calibration errors. Indeed, higher SNR may increase confidence in biased estimates when inversion efficiency is assumed rather than measured (35).

Compounding these hardware effects, labelling efficiency has emerged as a common pitfall. Our analysis shows that studies using the PASL labelling strategy FAIR sometimes report higher CBF estimates than those using pCASL. Indeed, FAIR models often assume a perfect inversion efficiency, which is historically why the technique has been favoured for its apparent robustness to B0 inhomogeneity (36). Empirical measurements in rodents, however, indicate that effective inversion efficiency is substantially lower and highly dependent on coil geometry, animal positioning, and magnetic field homogeneity (37). As a result, overestimating the labelling efficiency in the kinetic model may lead to a proportional overestimation of the required scaling. Thus, FAIR appears to be intrinsically more vulnerable to quantification bias (36, 38).

Using a PASL (FAIR) approach in rats, it was demonstrated that the labelling efficiency displayed marked sensitivity to the RF bandwidth of the inversion pulse in a rat model at 9.4T (39). This study showed that a bandwidth pulse greater than 15 kHz was required to robustly overcome the field inhomogeneity in the labelling region (39). Insufficient labelling can occur due the position of the volume coil in relation to the rat’s heart or the size of the transmit receive coil (40).

In contrast, pCASL explicitly exposes the vulnerability of labelling efficiency to off-resonance effects and vascular geometry, particularly at ultra-high fields (20, 21). While this increases technical complexity, it also encourages explicit calibration of inversion efficiency and closer inspection of labelling stability. Using a ramp to position the animal in the imaging sled can significantly reduce the variation of the labelling efficiency (23). The present findings therefore do not imply that pCASL is inherently more accurate, but rather that FAIR’s apparent robustness masks unquantified sources of bias that inflate perfusion estimates (21).

The ASL data acquisition and analysis approach vary considerably across each site for many reasons including scan time available to address the CBF research question in the study. The number of inversion times/ PLDs collected can affect the accuracy of CBF measurement. Also, the acquisition of separate M0, T1 and inversion efficiency maps will have an impact on the scaling factor used for the quantification of the CBF. This highlights the need for the preclinical community to come together to give recommendations for ASL protocols for robust CBF estimates in the rodent brain. In the interest of scientific transparency we are reporting our findings and attach all source data for others to consider further analysis.

Among biological factors, anaesthesia exerts a dominant influence on measured CBF (32, 41). Our *de novo* data clearly demonstrate the vasodilatory effects of isoflurane in producing higher baseline perfusion than vasoconstrictive agents like medetomidine, in agreement with prior studies (33). At the inter-site level, the lack of clear separation between isoflurane-only and mixed anaesthetic regimes likely reflects insufficient physiological reporting. Indeed, isoflurane concentration, ventilation status, and arterial CO_2_ tension are rarely reported with sufficient detail to allow normalisation across studies.

If animals are unventilated, respiratory depression under anaesthesia may cause respiratory acidosis due to a build-up of CO_2_ in the bloodstream, which is known to increase CBF by 100–200% (42, 43). Under extreme hypercapnia *PaCO_2_* >90 mmHg, CBF values exceeding 300 ml/100g/min have been reported and validated against autoradiography (43). More recently, CBF values exceeding 250 ml/100/min were reported in anaesthetised ventilated rats using 10% hypercapnia (3). Within this framework, the upper tail of CBF values observed in the metadata becomes physiologically plausible but not representative of normocapnic conditions. Yet without combined measurement of blood gases, such effects are difficult to disentangle from disease-related changes. Transcutaneous blood gas analysis offers a non-invasive way to monitor expired CO_2_ in mice (44). Since ventilation and blood gas analysis is impractical for longitudinal mouse studies a high breathing rate should be favoured by researchers concerned by unintended hypercapnia.

Finally, even under tightly controlled conditions, significant strain-dependent differences in baseline CBF were observed here in mice (Figure 3C) and have been previously reported in rats (21). These findings are in agreement with reports of anatomical variability in the circle of Willis and arterial branching patterns across rodent strains (45), but such differences are rarely accounted for explicitly in preclinical ASL studies. This suggests that some changes in baseline CBF attributed to pathology may instead reflect interactions between genetic background and vascular anatomy.

### Establishing a Normative Physiological Range

A central goal of this work was to assess whether a meaningful normative range for rodent CBF can be inferred from existing data. When unstratified, the literature suggests a broad range. However, when constrained by autoradiographic validation and physiological plausibility, a narrower range emerges (22, 43).

For mice under anaesthesia, the cortical CBF values average is 196 ml/100g/min when all the data studied are pooled. Whether it is analytically useful to pool so much data from different origins is left to the reader’s discretion but can serve as a reference point to those setting up rodent CBF studies. Values substantially above this value likely reflect hypercapnia, vasodilation, or quantification bias. Lower values may arise from vasoconstrictive anaesthesia, hypotension, or inefficient labelling. This average should be interpreted as a physiological anchor rather than a strict reference interval and deviations from it should prompt scrutiny of physiological state and ASL calibration.

## Conclusion

Our findings highlight that ASL-derived CBF is sensitive to both vascular physiology and modelling assumptions, as expected. Without rigorous control of these factors, apparent perfusion differences risk being misattributed to disease processes. To minimise the sources of bias, inversion efficiency should be measured whenever feasible, field-strength-specific relaxation constants must be used consistently and physiological monitoring should extend beyond respiratory rate to include ventilation or end-tidal CO_2_ where possible.

The trajectory of human ASL demonstrates that community guidelines dramatically improve reproducibility (10, 16). A similar effort in the preclinical domain would substantially improve translational alignment. Within the broader framework of this meta-analysis, these results reinforce that perfusion measurements reflect water mobility across vascular and tissue compartments only when physiological and methodological confounds are constrained. Careful attention to the pitfalls outlined above may help to narrow inter site variability which will benefit funders and other stakeholders.

## Supplementary materials

Appendix of Metadata *-*

Word file *Source data* Excel file

## Acknowledgements

This work was supported by a Wellcome Trust Career Development Award (I.N.C 306193/Z/23/Z CDA) and a Wellcome Trust Accelerator Award (Y.O. 316339/Z/24/Z), Grant LARSyS (FCT, DOI: 10.54499/LA/P/0083/2020, 10.54499/UIDP/50009/2020). We wish to acknowledge all authors affiliated with the meta-data, some of whom are listed here: Mark Lythgoe, Jack Wells, Constatino Iadecola, Laibaik Park, Louise van der Weerd, Leon Munting, Alexander Gourine, Patrick Hosford, Zhiliang Wei, Jan Klohs, Diana Kindler, Anusha Mishra, Zhenzhou Li, Benjamin Zimmerman, Emmanuel Barbier, Lydiane Hirshler, Philip Zhe Sun, Charith Pererea, Andrada Ianuș and Xavier Golay.

## Authors contributions

I.N.C conceived and directed the project; I.N.C, N.S. and P.F. designed the research; S.P.M., D.D., Y.O. and I.N.C. performed the research; S.P.M., D.D, Y.O. and I.N.C. analysed the results; S.P.M, Y.O. and I.N.C wrote the paper; All authors revised the article critically for important intellectual content.

## References

1. Buxton RB. Thermodynamic limitations on brain oxygen metabolism: physiological implications*. The Journal of Physiology. 2024;602(4):683–712.

2. Christie IN. Astrocytes: Orchestrators of brain gas exchange and oxygen homeostasis. The Journal of Physiology. 2025;n/a(n/a).

3. Hosford PS, Wells JA, Nizari S, Christie IN, Theparambil SM, Castro PA, et al. CO2 signaling mediates neurovascular coupling in the cerebral cortex. Nature Communications. 2022;13(1):2125–.

4. Munting LP, Derieppe M, Suidgeest E, Hirschler L, van Osch MJP, Denis de Senneville B, et al. Cerebral blood flow and cerebrovascular reactivity are preserved in a mouse model of cerebral microvascular amyloidosis. eLife. 2021;10:e61279–e.

5. Korte N, Barkaway A, Wells J, Freitas F, Sethi H, Andrews SP, et al. Inhibiting Ca2+ channels in Alzheimer’s disease model mice relaxes pericytes, improves cerebral blood flow and reduces immune cell stalling and hypoxia. Nature Neuroscience. 2024.

6. Buxton RB, Frank LR, Wong EC, Siewert B, Warach S, Edelman RR. A general kinetic model for quantitative perfusion imaging with arterial spin labeling. Magnetic Resonance in Medicine. 1998;40(3):383–96.

7. Paschoal AM, Woods JG, Pinto J, Bron EE, Petr J, Kennedy McConnell FA, et al. Reproducibility of arterial spin labeling cerebral blood flow image processing: A report of the ISMRM open science initiative for perfusion imaging (OSIPI) and the ASL MRI challenge. Magnetic Resonance in Medicine. 2024;92(2):836–52.

8. Pinto J, Chappell MA, Okell TW, Mezue M, Segerdahl AR, Tracey I, et al. Calibration of arterial spin labeling data—potential pitfalls in post-processing. Magnetic Resonance in Medicine. 2020;83(4):1222–34.

9. Pires Monteiro S, Pinto J, Chappell MA, Fouto A, Baptista MV, Vilela P, et al. Brain perfusion imaging by multi-delay arterial spin labeling: Impact of modeling dispersion and interaction with denoising strategies and pathology. Magnetic Resonance in Medicine. 2023;90(5):1889–904.

10. Alsop DC, Detre JA, Golay X, Günther M, Hendrikse J, Hernandez-Garcia L, et al. Recommended implementation of arterial spin-labeled perfusion MRI for clinical applications: A consensus of the ISMRM perfusion study group and the European consortium for ASL in dementia. Magnetic Resonance in Medicine. 2015;73(1):102–16.

11. Lindner T, Bolar DS, Achten E, Barkhof F, Bastos-Leite AJ, Detre JA, et al. Current state and guidance on arterial spin labeling perfusion MRI in clinical neuroimaging. Magnetic Resonance in Medicine. 2023;89(5):2024–47.

12. Qin Q, Alsop DC, Bolar DS, Hernandez-Garcia L, Meakin J, Liu D, et al. Velocity-selective arterial spin labeling perfusion MRI: A review of the state of the art and recommendations for clinical implementation. Magnetic Resonance in Medicine. 2022;88(4):1528–47.

13. Hernandez-Garcia L, Aramendía-Vidaurreta V, Bolar DS, Dai W, Fernández-Seara MA, Guo J, et al. Recent Technical Developments in ASL: A Review of the State of the Art. Magnetic Resonance in Medicine. 2022;88(5):2021–42.

14. Taso M, Aramendía-Vidaurreta V, Englund EK, Francis S, Franklin S, Madhuranthakam AJ, et al. Update on state-of-the-art for arterial spin labeling (ASL) human perfusion imaging outside of the brain. Magnetic Resonance in Medicine. 2023;89(5):1754–76.

15. Suzuki Y, Clement P, Dai W, Dolui S, Fernández-Seara MA, Lindner T, et al. ASL lexicon and reporting recommendations: A consensus report from the ISMRM Open Science Initiative for Perfusion Imaging (OSIPI). Magnetic Resonance in Medicine. 2024;91(5):1743–60.

16. Woods JG, Achten E, Asllani I, Bolar DS, Dai W, Detre JA, et al. Recommendations for quantitative cerebral perfusion MRI using multi-timepoint arterial spin labeling: Acquisition, quantification, and clinical applications. Magnetic Resonance in Medicine. 2024;92(2):469–95.

17. Fisher EMC, Bannerman DM. Mouse models of neurodegeneration: Know your question, know your mouse. Science Translational Medicine. 2019;11(493):eaaq1818.

18. Anderle S, Dixon M, Quintela-Lopez T, Sideris-Lampretsas G, Attwell D. The vascular contribution to cognitive decline in ageing and dementia. Nature Reviews Neuroscience. 2025;26(10):591–606.

19. Pell GS, Thomas DL, Lythgoe MF, Calamante F, Howseman AM, Gadian DG, et al. Implementation of quantitative FAIR perfusion imaging with a short repetition time in time-course studies. Magnetic Resonance in Medicine. 1999;41(4):829–40.

20. Hirschler L, Munting LP, Khmelinskii A, Teeuwisse WM, Suidgeest E, Warnking JM, et al. Transit time mapping in the mouse brain using time-encoded pCASL. NMR in Biomedicine. 2018;31(2):e3855–e.

21. Larkin JR, Simard MA, Khrapitchev AA, Meakin JA, Okell TW, Craig M, et al. Quantitative blood flow measurement in rat brain with multiphase arterial spin labelling magnetic resonance imaging. Journal of Cerebral Blood Flow & Metabolism. 2018;39(8):1557–69.

22. Buck J, Larkin JR, Simard MA, Khrapitchev AA, Chappell MA, Sibson NR. Sensitivity of Multiphase Pseudocontinuous Arterial Spin Labelling (MP pCASL) Magnetic Resonance Imaging for Measuring Brain and Tumour Blood Flow in Mice. Contrast Media & Molecular Imaging. 2018;2018(1):4580919.

23. Pires Monteiro S, Hirschler L, Barbier EL, Figueiredo P, Shemesh N. High-resolution perfusion imaging in rodents using pCASL at 9.4 T. NMR in Biomedicine. 2025;38(1):e5288.

24. Kozberg MG, Munting LP, Hanlin LH, Auger CA, van den Berg ML, Denis de Senneville B, et al. Vasomotion loss precedes impaired cerebrovascular reactivity and microbleeds in cerebral amyloid angiopathy. Brain Communications. 2025;7(3):fcaf186–fcaf.

25. Ratering D, Baltes C, Nordmeyer-Massner J, Marek D, Rudin M. Performance of a 200-MHz cryogenic RF probe designed for MRI and MRS of the murine brain. Magnetic Resonance in Medicine. 2008;59(6):1440–7.

26. Willie CK, Tzeng Y-C, Fisher JA, Ainslie PN. Integrative regulation of human brain blood flow. The Journal of Physiology. 2014;592(5):841–59.

27. Ramos-Cabrer P, Weber R, Wiedermann D, Hoehn M. Continuous noninvasive monitoring of transcutaneous blood gases for a stable and persistent BOLD contrast in fMRI studies in the rat. NMR in Biomedicine. 2005;18(7):440–6.

28. Kalthoff D, Po C, Wiedermann D, Hoehn M. Reliability and spatial specificity of rat brain sensorimotor functional connectivity networks are superior under sedation compared with general anesthesia. NMR in Biomedicine. 2013;26(6):638–50.

29. Patlak CS, Blasberg RG. Graphical Evaluation of Blood-to-Brain Transfer Constants from Multiple-Time Uptake Data. Generalizations. Journal of Cerebral Blood Flow & Metabolism. 1985;5(4):584–90.

30. Dobre MC, Uğurbil K, Marjanska M. Determination of blood longitudinal relaxation time (T1) at high magnetic field strengths. Magnetic Resonance Imaging. 2007;25(5):733–5.

31. de Vries RBM, Hooijmans CR, Langendam MW, van Luijk J, Leenaars M, Ritskes-Hoitinga M, et al. A protocol format for the preparation, registration and publication of systematic reviews of animal intervention studies. Evidence-based Preclinical Medicine. 2015;2(1):e00007–e.

32. Austin VC, Blamire AM, Allers KA, Sharp T, Styles P, Matthews PM, et al. Confounding effects of anesthesia on functional activation in rodent brain: a study of halothane and α-chloralose anesthesia. NeuroImage. 2005;24(1):92–100.

33. Munting LP, Derieppe MPP, Suidgeest E, Denis de Senneville B, Wells JA, van der Weerd L. Influence of different isoflurane anesthesia protocols on murine cerebral hemodynamics measured with pseudo-continuous arterial spin labeling. NMR in Biomedicine. 2019;32(8):e4105–e.

34. Brassard P, Labrecque L, Smirl JD, Tymko MM, Caldwell HG, Hoiland RL, et al. Losing the dogmatic view of cerebral autoregulation. Physiological Reports. 2021;9(15):e14982–e.

35. Msayib Y, Craig M, Simard MA, Larkin JR, Shin DD, Liu TT, et al. Robust estimation of quantitative perfusion from multi-phase pseudo-continuous arterial spin labeling. Magnetic Resonance in Medicine. 2020;83(3):815–29.

36. Kim S-G. Quantification of relative cerebral blood flow change by flow-sensitive alternating inversion recovery (FAIR) technique: Application to functional mapping. Magnetic Resonance in Medicine. 1995;34(3):293–301.

37. Muir ER, Shen Q, Duong TQ. Cerebral blood flow MRI in mice using the cardiac-spin-labeling technique. Magnetic Resonance in Medicine. 2008;60(3):744–8.

38. Duhamel G, Callot V, Tachrount M, Alsop DC, Cozzone PJ. Pseudo-continuous arterial spin labeling at very high magnetic field (11.75 T) for high-resolution mouse brain perfusion imaging. Magnetic Resonance in Medicine. 2012;67(5):1225–36.

39. Wells JA, Siow B, Lythgoe MF, Thomas DL. The importance of RF bandwidth for effective tagging in pulsed arterial spin labeling MRI at 9.4T. NMR in Biomedicine. 2012;25(10):1139–43.

40. Li Z, McConnell HL, Stackhouse TL, Pike MM, Zhang W, Mishra A. Increased 20-HETE Signaling Suppresses Capillary Neurovascular Coupling After Ischemic Stroke in Regions Beyond the Infarct. Frontiers in Cellular Neuroscience. 2021;Volume 15 - 2021.

41. Masamoto K, Kanno I. Anesthesia and the Quantitative Evaluation of Neurovascular Coupling. Journal of Cerebral Blood Flow & Metabolism. 2012;32(7):1233–47.

42. Sicard K, Shen Q, Brevard ME, Sullivan R, Ferris CF, King JA, et al. Regional Cerebral Blood Flow and BOLD Responses in Conscious and Anesthetized Rats under Basal and Hypercapnic Conditions: Implications for Functional MRI Studies. Journal of Cerebral Blood Flow & Metabolism. 2003;23(4):472–81.

43. Tsekos NV, Zhang F, Merkle H, Nagayama M, Ladecola C, Kim S-G. Quantitative measurements of cerebral blood flow in rats using the FAIR technique: Correlation with previous lodoantipyrine autoradiographic studies. Magnetic Resonance in Medicine. 1998;39(4):564–73.

44. Weber R, Ramos-Cabrer P, Wiedermann D, van Camp N, Hoehn M. A fully noninvasive and robust experimental protocol for longitudinal fMRI studies in the rat. NeuroImage. 2006;29(4):1303–10.

45. Ward R, Collins RL, Tanguay G, Miceli D. A quantitative study of cerebrovascular variation in inbred mice. J Anat. 1990;173:87–95.

